# Dynamic view of an allosteric intermediate in a positively cooperative dimer

**DOI:** 10.64898/2026.08.11.744235

**Authors:** Paul J. Sapienza, Trevor R. Mileur, Muhammad S. Khan, Kelin Li, Jeffrey Aubé, Andrew L. Lee

**Affiliations:** Division of Chemical Biology and Medicinal Chemistry, UNC Eshelman School of Pharmacy, University of North Carolina at Chapel Hill, Chapel Hill, NC 27516, United States; Department of Biochemistry and Biophysics, School of Medicine, University of North Carolina at Chapel Hill, Chapel Hill, North Carolina, USA

**Keywords:** Allostery, Entropy, NMR, Protein Dynamics

## Abstract

Positive cooperativity in ligand binding is a hallmark of allosteric oligomers, yet how the first binding event enhances the second remains obscure, because the pivotal singly-bound intermediate (lig₁) is thermodynamically disfavored and rarely accumulates. Distinguishing concerted (MWC) from sequential (KNF) mechanisms turns on one question: when a ligand binds one protomer, does its empty partner change conformation? Here we resolve this for the allosteric homodimer chorismate mutase (CM) using mixed-labeled dimers (MLDs), in which a single maleimide crosslink stabilizes a heterodimer carrying one NMR-labeled and one active-site-inactivated subunit, trapping lig₁ for prolonged study. Isothermal titration calorimetry shows that inhibitor binding to CM is positively cooperative and entirely entropy-driven, with the second event carrying far larger enthalpic and entropic swings than the first. Protomer-resolved NMR reveals that the first binding event switches both subunits to the relaxed (R) state—a concerted, MWC-like transition that rules out a strictly sequential model—yet the empty subunit is not a clean R conformer but a “fuzzy”, dynamically heterogeneous ensemble, with extensive microsecond– millisecond motion focused at the dimer interface, and a raft of residues surrounding the empty active site. Backbone probes confirm that both critical 11-12 loops adopt their active posture upon first ligand binding. The mismatch between the chemical-shift picture (MWC-like) and the thermodynamics (weighted toward the second event) argues that cooperativity is not encoded by a simple two-state switch, but by activated dynamics that a purely structural model cannot capture.

**Significance Statement:** Cooperative ligand binding underlies allosteric control across biology, but mechanisms have been hard to pin down because the crucial half-bound intermediate of a cooperative dimer barely exists at equilibrium. We stabilize this intermediate in the enzyme chorismate mutase by chemically linking one NMR-visible subunit to one that cannot bind ligand, letting us watch each subunit independently. Binding the first ligand flips both subunits to the active state, yet the empty subunit becomes highly dynamic rather than rigidly active. Combined with calorimetry, this shows that the free energy of cooperativity must be stored in ensemble dynamics, not merely in a switch between two structures. The linkage method further opens the door to additional solution-based studies required to better understand allostery and benchmark tools of the future.

## Introduction

Allostery is so central to the orchestration of biological processes that it was deemed “the second secret of life” half a century ago (1, 2). At a time when experimental determination of protein structures is giving way to their computational prediction (3), allostery remains an experimental frontier (4). Through investigations of structural and dynamic mechanisms of allostery (5, 6), design of allosteric proteins (7–12), and discovery of allosteric drugs (13–15), a wide range of research is looking to allostery to advance fundamental understanding of proteins and leverage it for new therapies and applications. The basic premise of allostery is that ligand binding or modification at one site of a protein affects activity or binding affinity at a distant site. Initially, the basis for allosteric cooperativity was conceived and understood in the protein class of homo-oligomers through either concerted ‘MWC’ (16) or sequential ‘KNF’ (17) conformational change of protein subunits. These subunit-level mechanisms became the dominant allostery paradigm, yet they also informed modern conceptions of monomeric protein allosteric activation like induced fit and conformational selection (18–20). The simplest case for testing these models is for an allosteric homodimer, in which discriminating between the two boils down to whether ligand binding in one protomer induces the conformational change (e.g., low affinity=’tense’=T to high affinity=’relaxed’=R) in its unbound partner. However, for a variety of reasons, examples of such a test – in homo-oligomers in general – are limited, especially in the case of positive ligand binding cooperativity since the intermediate liganded species (e.g., “lig_1_”) is thermodynamically disfavored. Such a direct, crucial test of allosteric mechanism thus remains elusive, and for the overwhelming majority of systems (21), it is unclear when, where, or if, allosteric protomers switch conformations.

An alternative, non-mutually exclusive, type of mechanism underlying protein allostery is that long-range cooperativity is mediated by dynamics, or modulation of dynamics (22–24). This is frequently discussed within a framework coined the “ensemble view of allostery” (5), and while it may have more obvious applicability to non-oligomeric proteins (e.g., monomeric), dynamics must also play an important role in classical, symmetric allosteric systems (25–28). It is interesting to note that the structure-based mechanisms (MWC or KNF T-R switching) for many classically allosteric systems are either unproven, unclear due to conflicting information between the crystal lattice and solution, or complicated (29–34). For these reasons, there is a lack of convergence in our understanding of allostery (4), or more practically, we lack the ability to predict ensembles and differences in allosteric energies. This is because the phenomenon is certainly more complex than the original models, and there is a need for greater detail and clarity of allosteric states, especially key singly-bound intermediate states in homodimers.

A largely-unexplored approach leverages NMR and mixed protomer labeling for capturing these elusive intermediates to answer significant questions in allosteric mechanism. In these dimers, one protomer is isotopically labeled and studied by NMR for its properties with the remainder of the oligomer unlabeled. We applied this approach first on *E. coli* thymidylate synthase (35), which has subtle allostery, but ultimately protomer exchange kinetics led to label scrambling. The solution is to crosslink protomers to stabilize the singly labeled species. A way to prevent subunit exchange, at least for homodimers, is to use bioorthogonal chemical groups installed at specific sites to link the labeled and unlabeled protomers. We achieved this using copper-click chemistry (CuAAC) on chorismate mutase (CM), as a proof of concept (36). While it can work well, we found the CuAAC strategy to be confounded by unpredictable yields and high cost. In this work we used a cysteine-maleimide strategy to construct “mixed labeled dimers” (MLDs) that isolate and stabilize the singly bound (or lig_1_) binding intermediate for the allosteric homodimer CM. Such MLDs enable prolonged study, via NMR, of the structural and dynamic features of the lig_1_ state, to consider MWC vs. KNF type behavior, as well as the dynamic properties.

Chorismate mutase (CM) from yeast is well-suited for careful studies of allosteric mechanisms because of its full complement of allosteric functions, small size for an allosteric enzyme, and excellent solution properties for NMR characterization. It catalyzes the conversion of chorismate to prephenate in the biosynthesis of tyrosine and phenylalanine. The homodimer binds two substrates in active sites >40 Å apart. The allosteric effectors tryptophan and tyrosine up- and down-regulate its activity (37) by binding to the effector binding region (EBR) (38), and in the absence of effectors its activity profile exhibits positive cooperativity, which is the focal point here. We use the CM transition state inhibitor (TSI) described by Bartlett (39) as a substrate analog ligand to capture the singly bound (lig_1_) intermediate for NMR observation. Isolation of pure lig_1_ is made possible upon appropriate active site mutation of one protomer, preventing formation of lig_2_. We show that wild-type CM binds TSI with a dramatic, entropically driven positive cooperativity. To get mechanistic insight into how binding of the first ligand increases the second by a factor of >10, MLDs were constructed to monitor both protomers of the lig_1_ state using either side-chain (^13^CH_3_-ILV methyl) or backbone (^15^N) NMR labeling. As shown previously, the TSI-bound protomer clearly adopts a stable R (“super-R”) conformation (40, 41). By contrast, the distal, unbound protomer does not adopt a single, stable conformation, as observed by broadened and missing NMR resonances. The chemical shifts, however, clearly agree with an R conformation over T, suggesting concertedness of the subunits. Hence, the dynamic signature of the empty lig_1_ protomer portrays a deviation from the assumption of intact T or R conformers and produces a new mechanistic picture of lig_1_ featuring a “fuzzy” empty protomer. The enhanced flexibility is not uniform; the ‘flickering’ of the empty subunit between states with distinct chemical shifts arises from loosening the dimer interface in both lig_1_ subunits around the region of a critical allosteric loop, releasing both loops from their inhibited ensembles to activated positions, and enhancing the flexibility of residues surrounding the empty active site. This straightforward approach paves the way for detailed structural and site-specific dynamics studies on CM and other dimeric systems that will be necessary to make allosteric prediction and design more routine.

## Results

### First and second active site binding events have dramatically different thermodynamic signatures

CM is positively cooperative in effector-free multiple turnover kinetics assays, requiring a Hill coefficient to fit the sigmoidal profiles (37). When trying to understand the mechanisms of cooperativity, this phenomenological parameter is limited because it conflates contributions from both *k*_cat_ and *K*_m_ and fails to separate the equilibrium and rate constants for the two binding and catalytic events, respectively. CM has been shown by us and others (37, 38) to have disparate K_m_ but similar *V*_max_ values among effector bound states—making it a K-type allosteric system. We therefore focused on the binding aspect of the interaction and measured the thermodynamics of a transition state inhibitor (TSI) binding to effector-free CM by isothermal titration calorimetry (ITC) (Figure 1A). The ITC isotherm was fitted using a two-site binding model with ΔH and ΔG (*K*_A_) defined separately for both binding events. The thermodynamics corresponding to the first and second binding events were very different, with the second, higher affinity binding exhibiting extreme changes in both ΔH and ΔS (Figure 1A). Both events are endothermic and driven by entropy, and the binding cooperativity is entirely entropy-driven, with -TΔΔS = −29.9 kcal/mol. In total this thermodynamic signature hints at a complete structural and/or dynamic remodeling of the system after the first ligand binding event. Specifically, the lig_1_ ensemble must look significantly different than lig_0_ (Figure 1B). However, for a positively cooperative dimer, the lig_1_ state does not accumulate significantly, making it difficult to study. We collected a methyl HMQC spectrum at the midpoint of a TSI titration to attempt to observe lig_1_ peaks directly. Lig_0_ and lig_2_ peaks are distinct (41), as expected for the conformational changes accompanying the T to super-R (sR) transition, and their resonance assignments are known. We used a binding polynomial to predict the fractions of species during a titration and, given the relative binding affinities reported here, the population of lig_1_ is expected to peak at ∼20% (Figure S1A), which in principle could be detectable by NMR. To classify the resonance positions of the lig_1_ species, we performed a simple TSI NMR titration, making sure to sample the lig_1_ apex condition. Interestingly this max [lig_1_] spectrum shows only two sets of peaks, each perfectly overlapping with either the lig_0_ or lig_2_ spectra (Figure S1B), showing that: 1) exchange between lig_0_ and lig_1Bound_ (and lig_1Empty_ and lig2) is slow on the NMR chemical shift timescale and 2) neither subunit of the lig_1_ species deviates significantly from the known T and R conformations to the extent that ILV resonances describe them. It is likely that binding of the first ligand induces the T to R transition in the local subunit because we see immediate buildup of bound-state intensity, thus lig_1B_ intensity overlaps with lig_2_, but whether the lig_1E_ intensity is subsumed by the lig_0_ or lig_2_ peak was not clear. Ultimately, site-specific NMR relaxation experiments to characterize the dynamics of the lig_1_ intermediate require resolved, uncontaminated peaks; and because each observed peak in the lig_1_-maximum spectrum contains contributions from multiple species, extraction of state-specific chemical shifts, line widths, and relaxation or exchange parameters is not feasible.

**Figure 1.**
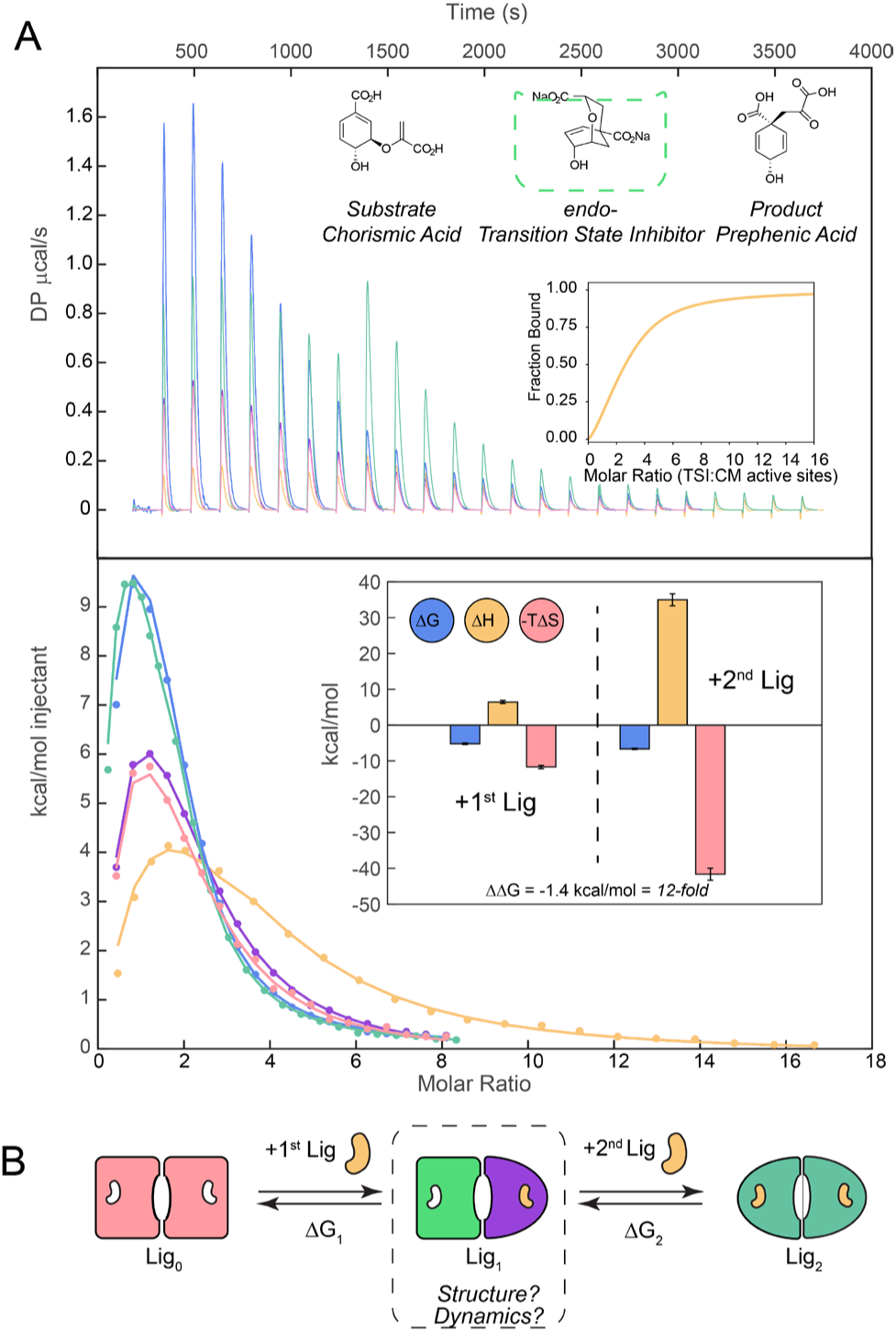
ITC analysis of TSI binding to effector-free CM. A) Global fit of five different c-values yields positive cooperativity. Top inset is overall fraction bound isotherm for one of the five experiments using the two fitted K_A_ values, bottom inset compares the underlying ΔG, ΔH, and TΔS thermodynamic parameters. B) Two different binding equilibria probed by ITC with the critical yet elusive intermediate lig_1_ state highlighted as the focus of this work.

### High-resolution observation of lig_1_ intermediate from mixed labeled dimers (MLDs)

To isolate the lig_1_ species and resolve signals from individual protomers, we prepared mixed-labeled dimers (35, 36) (MLDs) of CM (Figure 2). The purpose of an MLD is two-fold: 1) create a heterodimer with one protomer’s active site knocked out via mutation, such that ligand can only bind one protomer, and 2) isotope-label only one protomer for NMR observation. Because CM has rapid protomer exchange kinetics, an added requirement is that the MLD protomers be chemically linked to preserve labeled-unlabeled pairs. We previously showed a general-case approach to linking protomers directly through copper-catalyzed azide-alkyne cycloaddition (CuAAC) using unnatural amino acid (UAA) side chains (36). However, this approach was limited by cost in the way of unnatural amino acids and variable yields, inconsistent reactivity of the azide click partner despite attempts to regenerate it (42), and the strict need to perform the click reaction in a glove box without O_2_ (36). For CM, we found that a linkage via bismaleimide with PEG spacer was superior in terms of low cost, speed, and high, reproducible yields.

**Figure 2.**
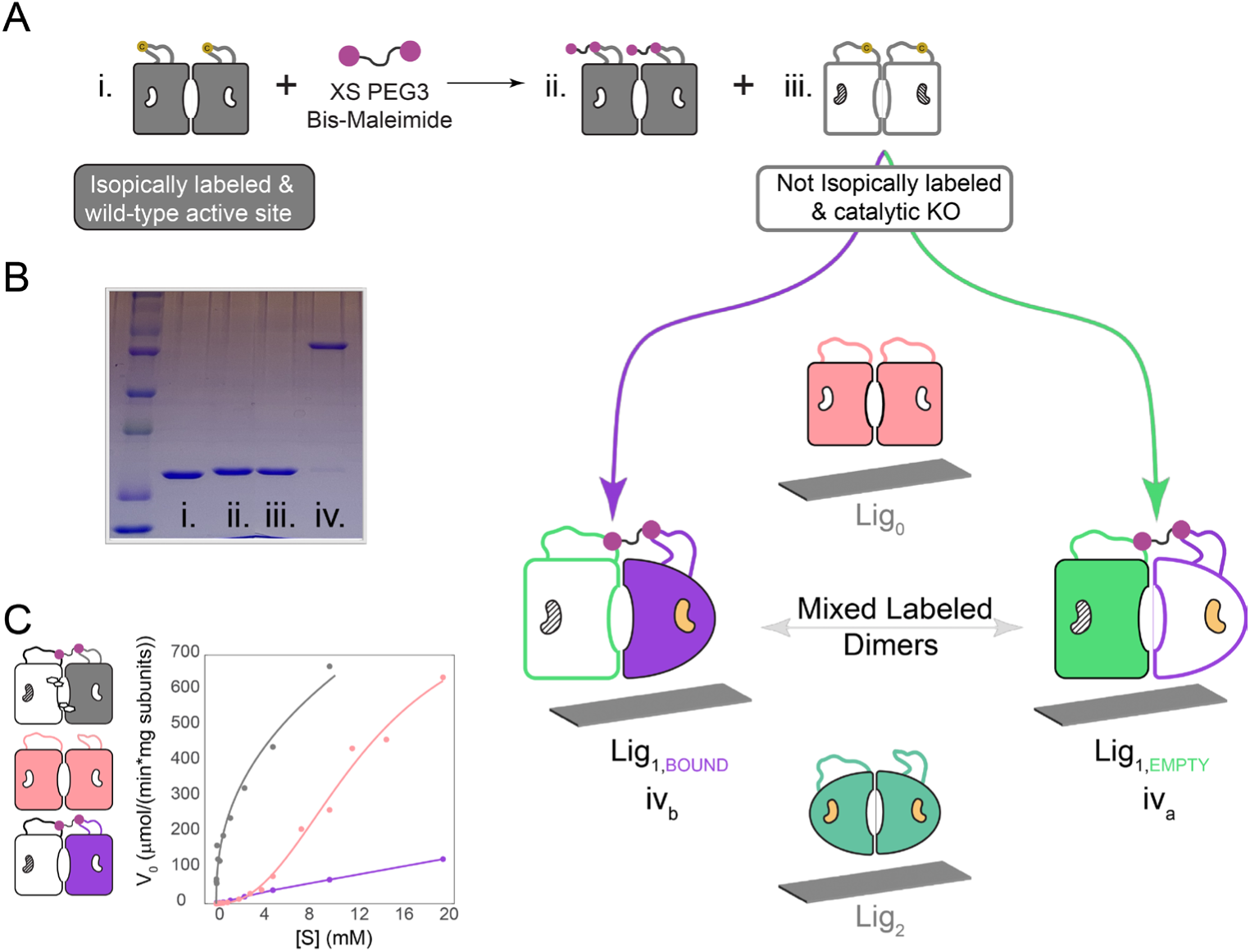
Bis-maleimide cross linking strategy for creating MLDs. (A) Sequential tagging, mixing, and cross-linking dimers to study lig_1_ empty and bound species. Solvent exposed cysteines are shown in yellow, filled subunits are isotopically labeled, and empty subunits are unlabeled. Hatched active sites are rendered unable to bind substrate. (B) Yield and purity are high. (C) MLDs give the expected single site, low affinity activity profile and retain the wild type feature of Trp activation.

Cysteines were installed in flexible loops having favorable inter-subunit geometry (Figure S2) and subjected to the workflow shown in Figure 2A. The expression yields are high without the need for UAAs, and the reactions are extremely efficient (Figure 2B). For samples to be used in NMR experiments, we were careful to use a slight molar excess of the unlabeled dimer in linkage reactions such that all labeled subunits are ‘soaked’ up. Indeed, MLD spectra showed remarkably little effect of linkage and no trace of the labeled homodimer (Figure S3A). MLDs retain the essential features of the wild-type enzyme in that they have similar NMR spectra (Figure S3A), are activated by Trp (Figure 2C), and the effector-free MLD with only one viable active site yields a right shifted and hyperbolic activity curve, consistent with the expected first low affinity binding event and removal of the positive cooperativity feature (Figure 2C). To show our findings are not influenced by, or idiosyncratic to the active site knockout mutations, we used two different pairs: R16L/N194A and R16L/T242G (Figure S3B). Both pairs abolish activity, and more importantly, neither shows any detectable TSI binding under NMR conditions (Figure S3C). In summary, the MLD technique gives us a unique handle on all three states responsible for the thermodynamics of allostery in a homodimer: The commonly studied lig_0_ and lig_2_ end states, and the two different MLDs, lig_1E_ and lig_1B_, as changes in either or both subunits can and likely do contribute to energetics of the elusive lig_1_ ensemble.

### The empty subunit of the lig_1_ state is in a dynamic R conformation

With the technology in hand, we set out to answer a fundamental question related to any cooperative homodimer: What is different about lig_1_ that modulates affinity of the second binding event relative to the first? To obtain the lig_1E_ methyl HMQC fingerprint we added a saturating amount of TSI to an MLD composed of an unlabeled wild-type active site subunit and an isotopically labeled R16L/N194A active site subunit (see Figure 2A, species iv_a_). It was immediately clear that the lig_1E_ state is highly dynamic on the μs-ms timescale based on severe line broadening relative to the lig_0_ and lig_2_ spectra (Figure 3A). The line broadening is a *bona fide* effect of distal active site binding because neither active site mutation, nor linkage manifests it in the apo form (Figure 3B). Further, dynamics do not result from bimolecular exchange that might be expected if the active site mutation weakened but did not completely abolish TSI binding (Figure S3C&D).

**Figure 3.**
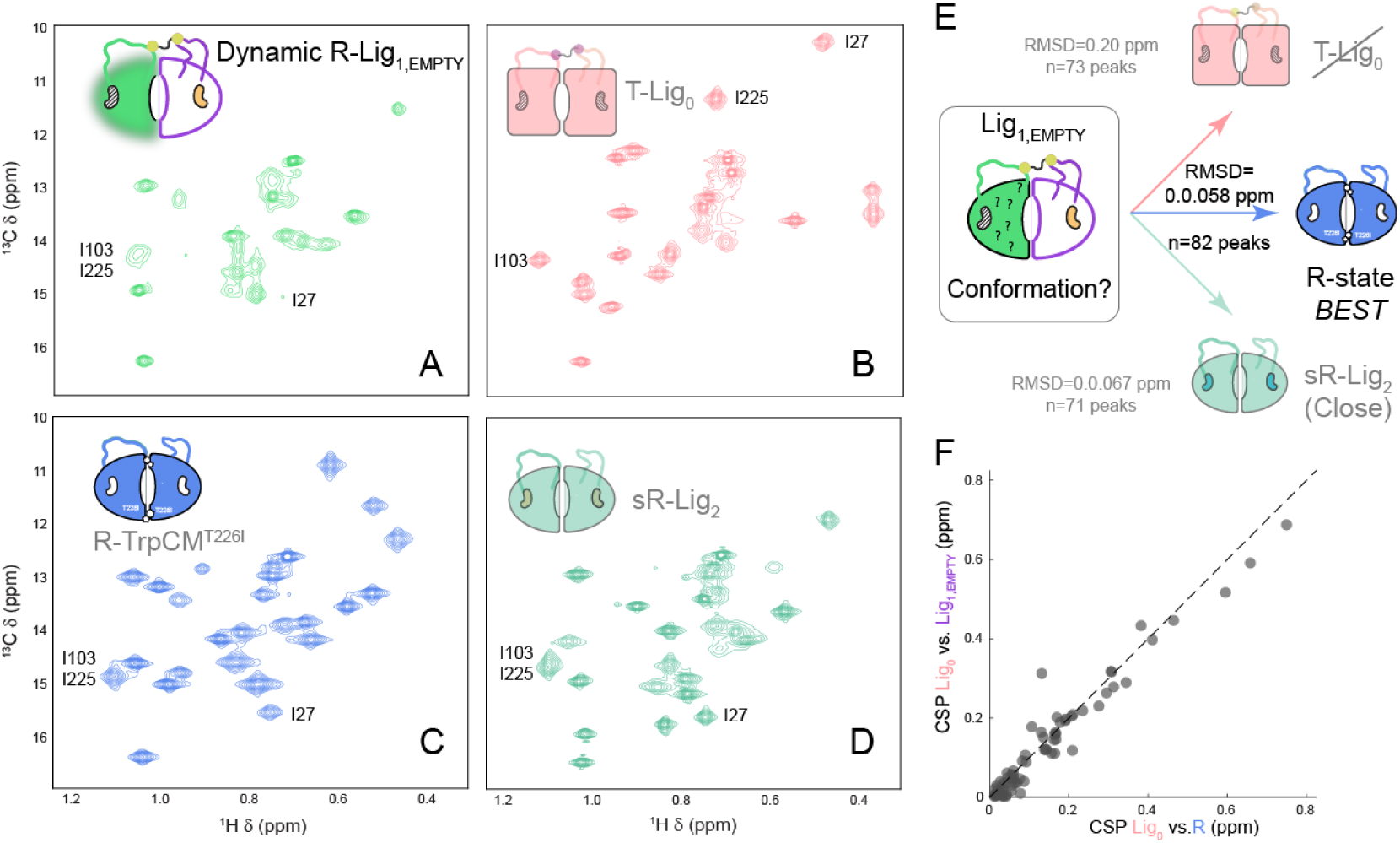
The empty lig_1_ subunit is a ‘fuzzy’ R-state. (A-E) Ile regions of ILV spectra are shown for clarity. Comparison of the lig_1E_ spectrum (A) with several other reference states: lig_0_-T conformation (B), lig_0_+Trp-R conformation (C), and lig_2_-super R conformation (D). Resonances in the lig_1E_ spectra are significantly broadened relative to the symmetrical reference states, revealing significant μs-ms dynamics in this intermediate. Several common peaks are shown in the four panels for comparison (E) ILV resonance RMSD analysis shows lig_1E_ most resembles the empty R-state. (F) Linear CSP analysis of all ILV peaks reveals essentially a global switch of the lig_1empty_ subunit to the R-state.

We then compared the peak positions of lig_1E_ with other assigned reference states: T, R, or sR (Figures 3B-D), and an RMSD analysis shows the empty subunit spectrum clusters with the R fingerprint (Figure 3E). The switch of the distal subunit from T to R is essentially global in nature as evidenced by the linear correlation of a set of 82 methyl CSPs (Figure 3F). It is noteworthy that the spectra of the active site-empty R-state and the active site-bound sR-states are highly similar (41), yet the match of lig_1E_ with the empty R-state is better than that of bound sR (Figure 3E). The RMSD is based on a heavily overlapping but not formally identical set of peaks. To rule out bias and to confirm that the switch is not an artifact of residual low affinity active site binding, we looked at two resonances, I192 and I239, whose peak positions are diagnostic of TSI in the active site, and these two are unequivocally in the empty active site positions in the lig_1E_ spectrum (Figure S4).

As stated above, binding of the first ligand activates widespread flexibility within the distal subunit. The most parsimonious explanation is the lig_1E_ R-like ground state samples the T conformation. To address this possibility, we plotted the lig_1E_ HMQC intensities, which are an inverse proxy for exchange, against the T vs. R change in chemical shift (Figure 4A). While there is a correlation—the resonances with the largest CSPs have among the lowest intensities and vice versa—more complex dynamics must be at play. We identified ∼30 resonances in the lower left quadrant of the plot that are significantly more broadened than what is expected from a simple R to T event. These residues map to the EBR (Figure 4B purple), and a raft of residues that travel from the dimer interface to a region that surrounds the empty active site (Figure 4B pink). We tested the generality of these findings by looking at a lig_1E_ species with a different active site mutant pair, and both the switch to R and increased dynamics are reproduced (Figure S5A) suggesting we are observing the wild-type allosteric repertoire. Taken together these data show binding of the first ligand causes the second subunit to adopt the R-conformation, one that ‘flickers’ between unknown conformational states, conferring but a dynamic ‘personality’ that is distinct from any symmetric state observed thus far. This behavior is somewhat reminiscent of the MWC model that mandates a concerted T to R switch, but with an additional layer of dynamic detail that can only be provided by a significant effort to trap the lig_1_ state, coupled with NMR spectroscopy.

**Figure 4.**
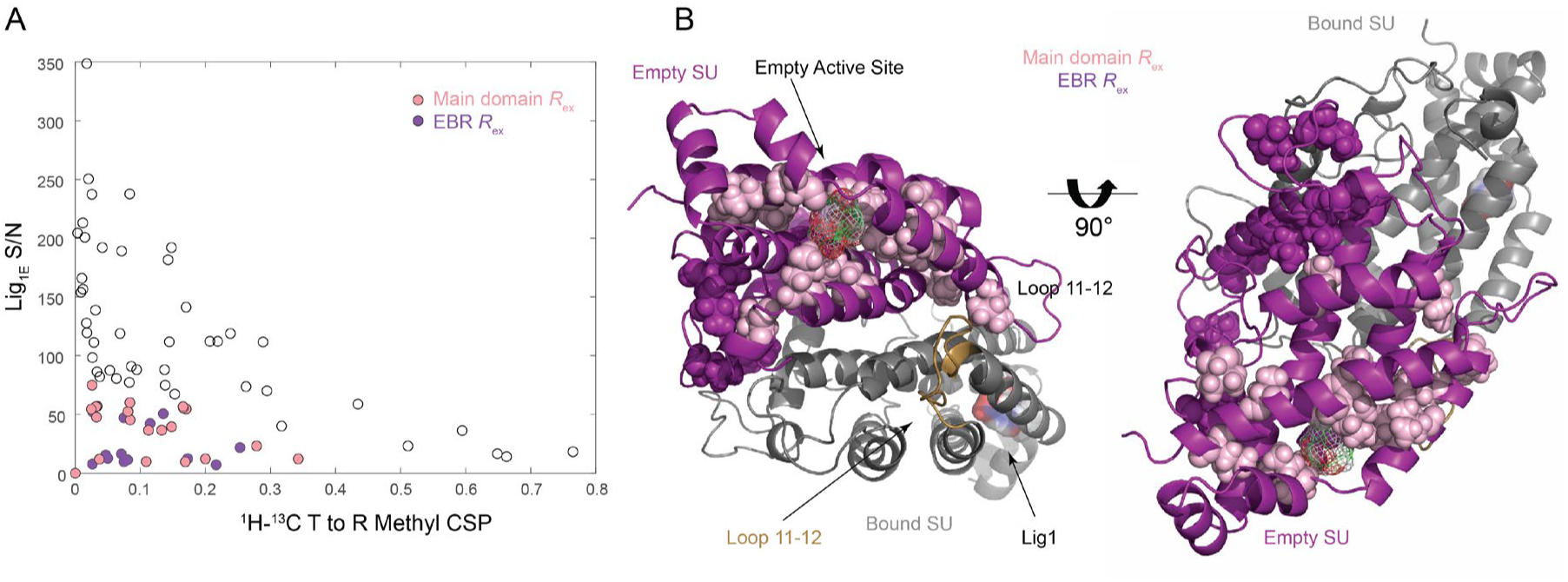
μs-ms dynamics in lig_1E_ map to the distal active site and EBR. (A) Lig_1E_ HMQC spectrum intensities are plotted against the chemical shift perturbations (CSPs) associated with the T to R conformational change. Despite the rough correlation, a large group of peaks in the lower left quadrant exhibit excess broadening beyond what is expected from R to T switching. These are highlighted in space fill on the structure (B) within the EBR (purple) and main domain (pink). Note how these dynamics spread from the dimer interface at the stubs of loop 11-12 to a cluster of residues that surround the empty active site shown as a wire framed TSI. Loop 11-12 is labeled in the empty subunit and shown as contrasting gold color in the bound.

### Ligand binding propagates asymmetrically between protomers

To examine the bound subunit of lig_1_, we constructed a reciprocal MLD in which the labeled subunit is able to bind TSI and the unlabeled partner is not (Figure 2A-species iv_b_). The methyl HMQC spectra of this MLD (lig_1B_) and that of lig_2_ overlay nearly perfectly (Figure 5A), indicating that binding to the first active site triggers what is essentially a complete switch to the super-R conformation in the local subunit. Ligand binding propagates differently depending on whether a subunit is being bound or sensing it from across the subunit interface, and this is only discernible with extreme negative cooperativity or MLDs. To appreciate the complete scope and scale of the chemical shift changes from the two binding events, we compared four chemical shift perturbation vectors derived from the peak positions of the two symmetrical end states and two MLDs (Figure 5B): local_1_ and distal_1_ report on how ligand one binding is sensed at the local and distal subunit, respectively, and local_2_ and distal_2_ describe how the second binding event is propagated. This perspective shows the dominant perturbations in both subunits are caused by the first binding event. Also, there are smaller, yet significant shifts in the empty lig_1_ subunit upon binding the second TSI molecule, and finally, the pre-bound lig_1_ subunit senses the second ligand binding event, but the shifts are subtle (Figure 5C&D). These subtle distal_2_ shifts map to the dimer interface and ‘stubs’ of loop 11-12 (Figure 5D). Although they are small, we believe these CSPs are indeed a signal, reporting on important features of the lig_1_ ensemble because they are reproduced in two different MLDs (Figure S5B) and cluster to a region near loop 11-12 (see below) and directly opposed to the core of dynamic residues in the empty subunit.

**Figure 5.**
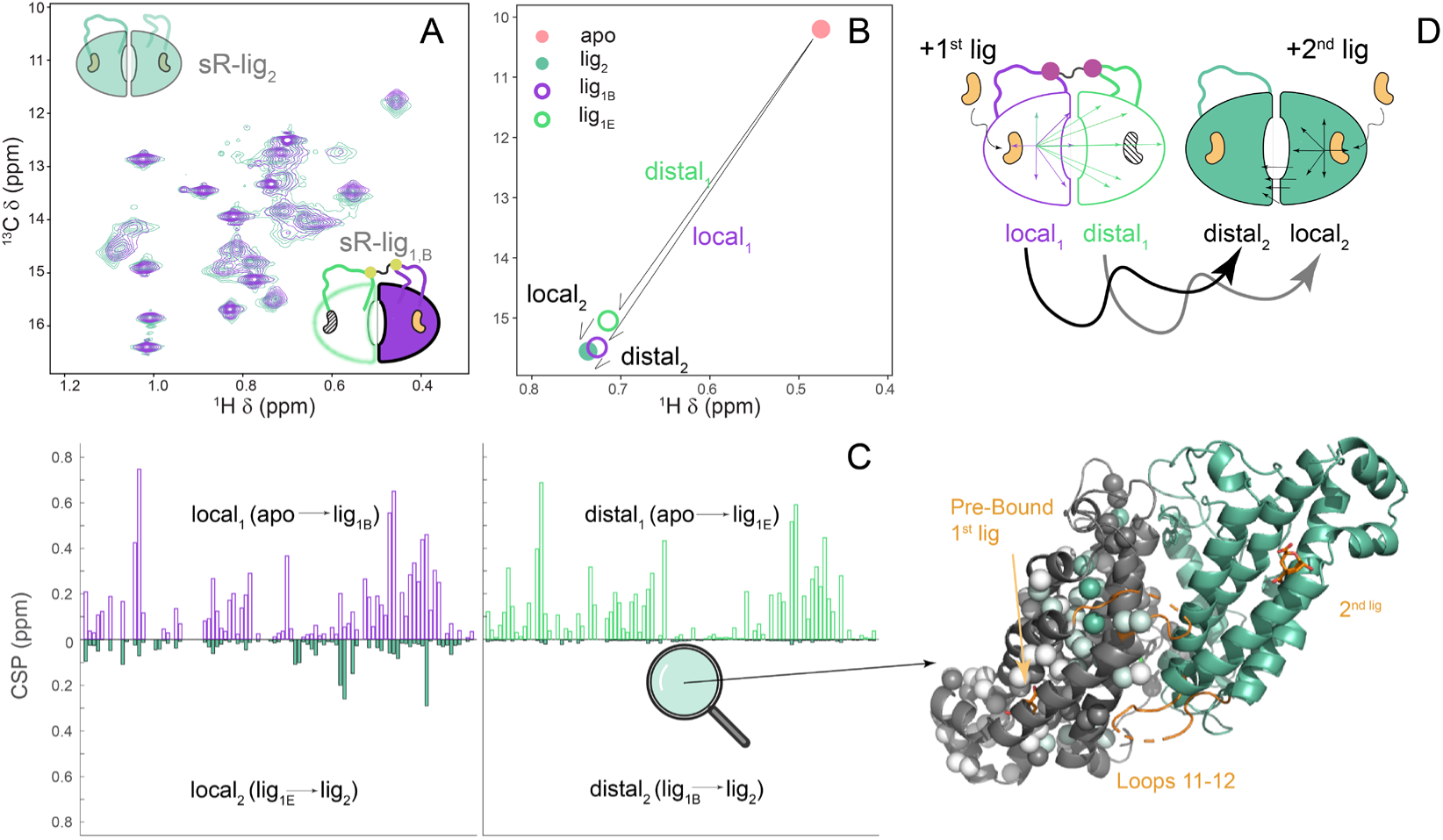
First binding event dominates chemical shift changes throughout both subunits. (A) Local subunit of lig_1_ looks remarkably like lig_2_. (B) CSP vector definitions report on how each subunit senses both binding events. (C-left) The bar charts plot these vectors to demonstrate that local_1_ and distal_1_ dominate the CSP, the empty subunit shifts somewhat upon binding the second ligand (local_2_), and the prebound subunit of lig_1_ only subtly adjusts peak position upon second ligand binding (distal_2_). (C-right) The distal_2_ CSPs that are too small to evaluate on the expanded common y-axis scale are magnified and painted on the prebound subunit. The changes cluster to the nexus of loops 11-12 and the dimer interface. (D) Cartoon summarizes the bar charts. First binding event has large local and distal subunit perturbations, whereas the second has modest effects on the empty subunit and localized minor effects on the pre-bound subunit.

### Lig_1_ binding triggers the active ensemble of a critical dynamic loop

Loop 11-12 requires special consideration in CM because several lines of evidence show it is central to allosteric regulation: i) loop chimeras can shuffle the regulatory features of the enzyme (43), ii) a single point mutant at the base of the loop was shown to constitutively activate CM (44) regardless of effector status, and iii) our previous work demonstrated that the charged flexible loop transiently approaches the active site and modulates the electrostatics throughout the protein in activator— but not inhibitor bound CM (Figure 6A) (45). This observation has the potential to resolve the paradox that activated TrpCM is in the T, not the R-state as its major conformer, and provides a mechanism for binding affinity enhancement independent of a T vs. R structural switch. However, the loop lacks ILV probes aside from I225 in the ‘stub’ (Figure 6), so methyl HMQC spectra are blind to whether the loops are coupled to the rest of the main domain in lig_1_. Backbone amides, in principle, yield NMR probes for all non-proline residues and loop chemical shifts are assigned and are distinct for the inactive and active conformations. We measured TROSY ^1^H-^15^N HSQCs for all three states and, strikingly, the resonances of both lig_1_ loops align with the R- and not the T-state (Figure 6B). Beyond loop 11-12, the collection of backbone spectra reinforces the conclusions drawn from methyl probes, in that both subunits switch to the R conformation upon binding of the first ligand—albeit dynamic in the empty protomer— with the added information that the change in 11-12 loop posture occurs simultaneously. The backbone and methyl data are also in accord with respect to showing that the distal subunit is ‘fuzzy’ based on widespread broadening and ∼30 missing peaks in the lig_1E_ HSQC (Figure S6). Further, analysis of a comparison of lig_1B_ spectrum intensities with those in lig_2_, show enhanced flexibility spreads beyond the empty subunit to several regions in the bound protomer: at the dimer interface where the stubs of loop 11-12 emerge from the bound subunit, and regions of helices 2 and 11 where loop 11-12 from the distal, empty subunit approaches (Figure 6C). Collectively, these data point to a model in which the first ligand binding event loosens the dimer interface in both subunits where the stubs of both loops 11-12 emerge. This frees the distal subunit to adopt an R conformation ensemble with active site flexibility that imparts higher affinity for lig_2_ (Figure 6D).

**Figure 6.**
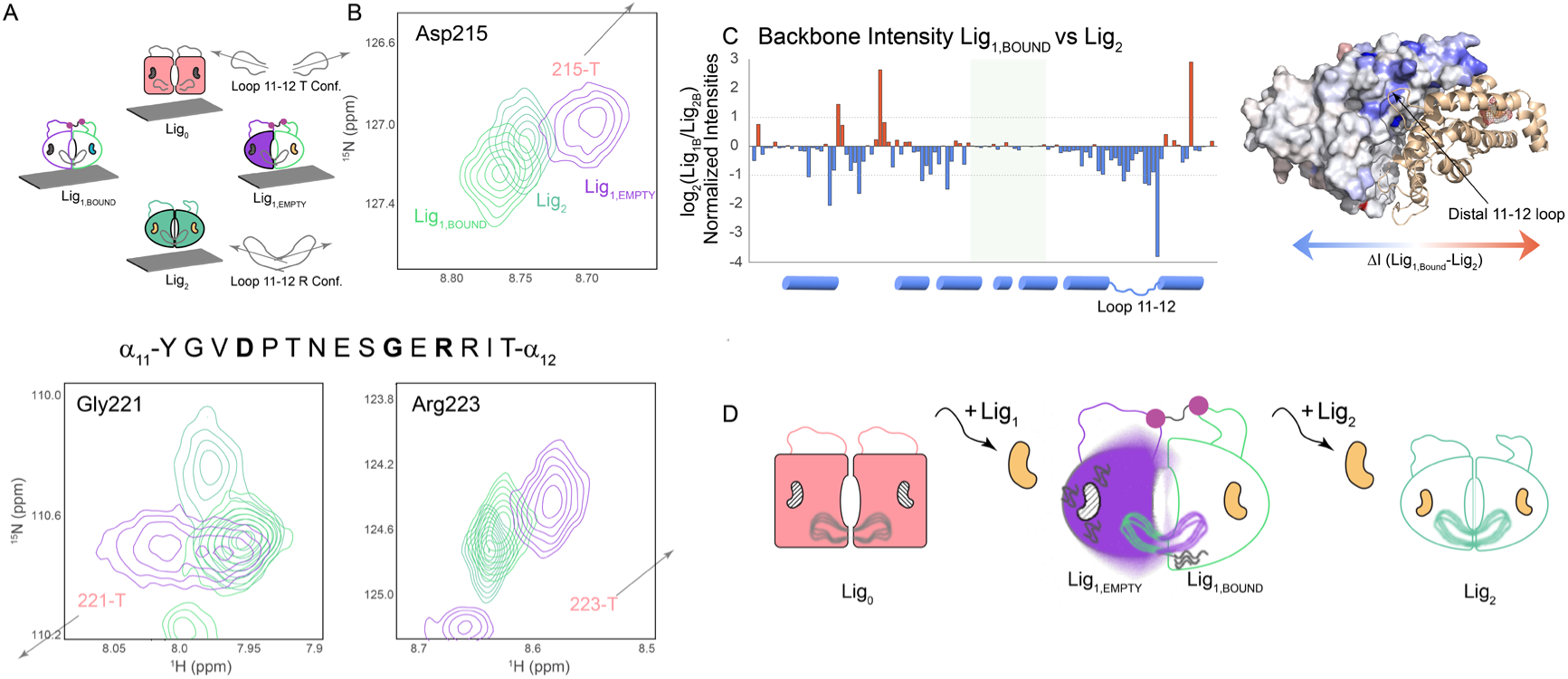
Loops 11-12 of both lig_1_ subunits adopt the active ensemble. (A) Schematic of the inactive and active poses of loop11-12 in which the loop is oriented within and across subunits, respectively. (B) Three representative loop 11-12 amide resonances having distinct active vs inactive chemical shifts match the lig_2_ position in both lig1_E_ and lig1_B_. (C) Comparison of lig_1B_ vs lig_2_ amide resonances show line broadening in lig_1B_ loop 11-12 and the region that senses loop 11-12 of the distal subunit. Prebound subunit is shown as a surface, residues with line broadening relative to the lig_2_ reference state are painted with a blue gradient. Spectra were collected at similar concentrations, number of scans, and receiver gain and were normalized by the residues in the greyed window. This group of residues shows little to no CSP and no *R*_ex_ in any of the states examined. (D) Integration of chemical shifts and linewidths from 4 states into a model in which the apo state is locked in a low dynamics T-state with respect to both the main domain and loop 11-12. The first binding event triggers a concerted switch of both subunits to R. However significant dynamics are activated in the distal subunit and the local subunit at the dimer interface nexus of both loops 11-12 that propagates to the empty active site.

## Discussion

Mixed labeled dimer technology has provided a rare glimpse into the nature of the key lig_1_ intermediate in an allosteric homodimer. Protomer specific NMR spectra of lig_1_ reveal a concerted switch of both CM subunits to the R-state. Taking the chemical shift coordinates as a proxy for structure might suggest CM adheres to the classic MWC model of allosteric mechanism. However, several lines of evidence support the view that this concerted switch may not be the sole or dominant driver of positive cooperativity. The first is our previous observation that the degree of CM activation by tryptophan is not correlated with the R-state population. Most notably we showed that a mutant of CM is fully activated with either Trp or Tyr bound, yet the first form is entirely in the R conformation and the second in T (46). We also uncovered a reciprocal case in which CM is in the R-state but has an inhibited activity profile (46). The second line of evidence is the thermodynamic signature of the two binding events shown here by ITC. Both are entropy-driven and enthalpically opposed (positive ΔH, favorable −TΔS), but the *second* event has a dramatically larger swing in both—a much bigger enthalpic penalty that is offset by a larger entropic gain—for a net ΔΔG of only ࢤ1.4 kcal/mol. If the concerted T→R shift is triggered by the first ligand (as our NMR data show), the bulk of that shift’s enthalpic/entropic signature should show up in ΔH_1_/ΔS_1_, not ΔH_2_/ΔS_2_. Yet, we see the opposite—the second step carries the larger thermodynamic weight—which is difficult to reconcile with a model in which the concerted structural shift provides the cooperative free energy. In other words, even in this case of average structure symmetry in lig_1_, the ensemble must be highly asymmetric in terms of structure and/or dynamics. We note that the ordering of the fuzzy subunit upon lig_2_ binding is naively expected to *cost* entropy, which is opposite to what we see. However, there is limited entropy— *Rln*(2) — stored in the type of coupled two-state exchange that leads to NMR line broadening; rather it is the distributed ps-ns dynamics measured experimentally by the NMR order parameter, *S*^2^, that relate to conformational entropy (47), and *R*_ex_ and *S*^2^ need not correlate (28). Thus, water release or activation of ps-ns dynamics is expected to provide the entropic driving force for the second binding event. Taken together, it is essential to look beyond the two-state conformational change to account for the massive swings in ΔH and TΔS, and CM provides another vivid example of the complexity of allostery, and the need to pivot to ensemble descriptions.

This highlights understanding and therefore designing allostery as exciting untraversed frontiers in biomedical science. Protein structure prediction has already crossed this boundary as tools built on deep learning, exemplified by AlphaFold3 (3) (AF3), now generate accurate atomic-resolution models, collapsing what used to be a long experimental campaign into a single computational query. However, AF would fail at predicting allostery in CM and many other systems because it is more complex than the T and R models deposited in the Protein Data Bank (PDB). We believe this gap might be especially salient in considering homodimers, the most abundant protein class in nature (48), in which allostery is governed by the difference in free energy between the first and second ligand binding events. To understand regulation at the most basic level in these systems, one would want to know the structure of the apo (lig_0_), lig_1_, and saturated forms of the dimer and compare the lig_1_ and lig_0_ models in search of features underpinning positive or negative cooperativity. Unfortunately, this is rarely achieved as evidenced by a sampling of 223 high resolution crystal structures of which only 11 are asymmetric (21). Thus, the AF3 training set is highly symmetrized, so any attempt to model an asymmetric lig_1_ complex with a tool trained on such a symmetric ‘diet’ must be viewed with healthy skepticism.

How might allosteric free energies and mechanisms ultimately be predicted with the ease at which we currently generate accurate static structures? The success of structure prediction largely rests on the existence of the PDB: hundreds of thousands of solved structures, assembled over decades into a single standardized resource. No comparable repository exists for the energetics and dynamics that underlie allostery. Structural, thermodynamic, and kinetic parameters (chemical shifts, NMR order parameters, binding free energies, exchange rates, populations of conformational substates) are scattered across the literature in disparate formats, system by system, each requiring a dedicated multidisciplinary campaign to obtain. The result is that while a structure can now be predicted in seconds, the free energy landscape that governs function has had to be measured experimentally, system by system. The Allosteric Database (49) has, since 2009, assembled a large, actively maintained catalog of allosteric proteins, sites, and modulators — the closest analog to a PDB-scale resource this field has — but it captures structural and pharmacological annotation, not the thermodynamic or dynamic parameters (ΔG, ΔH, ΔS, exchange rates, substates and populations) that actually define an allosteric mechanism. Emerging generative approaches — ensemble predictors like BioEmu (50), a recent incarnation of AF3 that generates ensembles guided by experimental data (51), and other workflows (52) — represent promising steps toward predicting conformational populations and perhaps ultimately their energies directly. However, the field stands at an early stage where computational predictions of allosteric structure and energetics need to be rigorously tested against quantitative experimental ground truth. MLDs offer an experimental handle, and CM offers an unusually well-characterized system for generating exactly this kind of benchmark: a symmetric homodimer with well-defined allosteric effectors, an established kinetic and thermodynamic framework spanning decades, and is amenable to NMR, which is the tool of choice to provide the necessary structural and dynamic allosteric information. Lastly, it is noteworthy that the NMR studies capturing and/or studying asymmetric lig_1_ states tend to be negatively cooperative (27, 53, 54), with a notable exception (55) where the chacteristics of lig_1_ were inferred by intensity analysis rather than isolating it, consistent with cases of positive cooperativity remaining a significant challenge

The lig₁ intermediate’s most distinctive feature — a “fuzzy,” dynamically heterogeneous empty subunit rather than a clean T or R conformer — raises questions our thermodynamic data and NMR fingerprints cannot yet resolve: 1) how much of the large −TΔS_₂_ term reflects newly liberated backbone and side-chain motion (configurational entropy) versus altered hydration? These two entropy sources are not distinguishable from ITC or chemical shift data alone. 2) what are the dynamic events that lead to the extensive line broadening in lig_1_? The degree of signal loss does not scale with the T to R chemical shift change so sampling the T state cannot be the sole μs-ms timescale dynamic event, and 3) are the activation mechanisms initiated by the first ligand binding event and Trp binding the same? It is interesting that the areas showing dynamics that deviate from clean T to R or R to T switches are the same in TrpCM and effector-free lig_1E_ CM, respectively (Figure 5 vs. ref. 41). In both cases, residues within the EBR and active site perimeter are dynamic outliers. And this, coupled with the fact that the active site is not solvent accessible in any of the deposited T, R, or super x-ray models, lead us to hypothesize that this flexibility is related to activation by both Trp and the first substrate binding event. The relaxation and exchange measurements already applied elsewhere in our work on this system — backbone and side-chain order parameters, CPMG (41), CEST (41), and PRE (45) — can be extended directly to the lig₁_B_ and lig₁_E_ MLDs and asymmetrical mixed forms of other effector states to answer these questions. More broadly, MLDs will facilitate similar investigations of the large class of allosteric oligomers.

## Materials and Methods

### Expression and purification of CM

Unlabeled, U-[^2^H,^15^N], U-[^2^H,^13^C, ^15^N] and {U-[^2^H,1^5^N]; Ileδ1-[^13^CH_3_]; Leu, Val-[^13^CH3,12CD3]} CM were expressed and purified as described (41) with the exception that isotopically labeled preparation used the M9+ formalism of Clore and coworkers (56) to make more efficient use of D_2_O. When ILV protein was expressed, 16 and 30 mg of the α-ketobutyric and α-ketoisovaleric acid precursors respectively were added per 100 ml culture. For methyl-methyl NOESY experiments, we used a different α-ketoisovaleric acid precursor to generate {U-[^2^H,^15^N]; Ileδ1-[^13^CH_3_]; Leu, Val-[^13^CH_3_,^13^CH_3_]} CM, which allowed NOEs between geminal methyl groups in the same LV residue. And for the ‘nearest-neighbor’ (57) ^13^C constant time HMQC to discriminate Leu resonances from Val, we used a U-[^2^H,^13^C,^15^N] media supplemented with uniformly ^13^C labeled α-keto precursors.

### Synthesis of Transition State Inhibitor

TSI was synthesized as described previously (36, 41) and obtained as a mixture of the 3-endo-8-exo and 3-exo-8-exo diastereomers. ITC thermograms acquired with this mixture in the syringe were sufficiently noisy that reliable fits were not possible; the flat baselines shown in Figure 1 were achieved only after isolation of the pure (1R,3S,5S,8R) enantiomer, described in detail in SI Appendix. The active enantiomer corresponds to the transition state inhibitor modeled in the super-R crystal structure of CM (PDB 3CSM) (40). NMR spectra acquired with the diastereomeric mixture and with the pure enantiomer are indistinguishable, and the (1S,3R,5R,8S) enantiomer showed no detectable binding by NMR; accordingly, ITC was performed with enantiopure TSI, while NMR samples used the mixture and were verified spectroscopically to be at saturation in every case.

### Isothermal Titration Calorimetry

CM was dialyzed into NMR buffer: 25 mM NaPO4, 150 mM NaCl, 1 mM EDTA, 0.5 mM TCEP, pH 6.5. TSI was resuspended in water and being a diacid, was titrated to pH 6.5, lyophilized, and resuspended in the dialysate above. The concentration of TSI was determined by NMR using PULCON (58) with L-Tyr and L-Trp as standards. Five experiments were conducted in a Malvern PEAQ ITC with [CM] in the cell at 28, 28, 55, 56, and 110 μM, paired with five different TSI syringe concentrations of 0.56, 1.12, 1.1, 2.24, and 2.2 mM to give five different C-values. Titrations were conducted at 15 °C. Thermograms were integrated using NITPIC (59) and were fit globally using the two site sequential model in SEDPHAT (60). The thermodynamic parameters derived from microscopic *K*_A_ values are reported and discussed herein.

### MLD Assembly

Constituent subunits were expressed and purified separately in reducing agent to ensure the reactivity of cysteines. Each enzyme was then exchanged into NMR buffer without reducing agent prior to linkage, which was accomplished in two steps. In step one, all subunits of the first dimer at 10 μM were tagged using a 20-fold excess bis-mal-PEG3 (BroadPharm) over subunits on ice, which prevented any trans dimer reactivity (Figure S7A) for temperature effects and Figure 2A for scheme and SDS gel). A Cys to Ser mutation was required at position 39 in the tagging subunit to prevent intra subunit crosslinking (Figure S7B). Unreacted tag was then removed with a Sephadex G-25 column and a stoichiometric amount of the second dimer was mixed in. Dimer subunits exchange readily and covalent linkage was complete within 15 minutes. We were careful to use a slight excess of the unlabeled dimer in the mixing step to ensure that only the MLD, and not the labeled homodimer would give signal in NMR experiments. A description of the mutants and MLDs used in this work are in Table S1.

### Backbone and ILV Resonance Assignments

The assignment strategy for several reference states was reported elsewhere (41). Analysis in this work was aided by the backbone resonance assignments of TyrCM and TSI-TrpCM, which allowed assignments for the first time of the entire loop11-12 in both T and R states. Briefly, we back exchanged U-[^2^H,^13^C, ^15^N] CM for 24 hours in 2M Urea NMR buffer, pH 9, removed the urea at pH 9 by dialysis, dialyzed against NMR buffer, added the appropriate ligands to saturation, and acquired the standard suite of TROSY HNCO, HN(CA)CO, HNCA, HN(CO)CA, HN(CA)CB, and HN(COCA)CB triple resonance experiments. Sequential connections were made using the NMRViewJ RUNABOUT assignment module. The MLD backbone assignments discussed herein were transferred from the array of assigned reference states. The ILV methyl assignments for ApoCM, TrpCM, TrpCM^226I^, and TSI-TrpCM were reported previously (41). We used a similar NOESY (61) and guided MAUS auto-assignment strategy (62) to assign ILV resonances in the binary TSI-CM complex. MAUS is aided by a priori knowledge of which resonances are from L or V, and this accomplished by a modified constant time HSQC as described (57). As with the backbone, MLD ILV peak assignments were transferred based on reference states aided by a single methyl-methyl NOESY (61) to help resolve some peak movements that resulted from the active site mutations. Weighted heteronuclear (^1^H-X) chemical shift perturbations were calculated with coefficients of 0.133 and 0.14 relative to proton for carbon and nitrogen respectively.

### NMR Spectroscopy

NMR data were acquired on a Bruker Avance III HD 850 MHz four channel spectrometer equipped with a TCI H-C/N-D 5 mm CryoProbe unless otherwise specified. Pulse sequences and parameter sets were from the Bruker library for HNCO (trhncogp3d, 50% non-uniform sampled), HN(CA)CO (trhncacogp2h3d, 600 MHz with TCI cryoprobe), HNCA (trhncagp2h3d2), HN(CO)CA (trhncocagp2h3d, 600 MHz), HN(CA)CB (trhncacbgp2h3d), HN(COCA)CB (trhncocacbgp2h3d, 600 MHz), and TROSY-HSQC (trosyf3gpphsi19.2). The pulse sequence for the nearest neighbor constant time HMQC was published (57), and the sofast 3D HMQC-NOESY-HMQC pulse program (61) was made available for download by the University of Minnesota NMR Center. Two NOESYs were acquired for binary TSI-CM: one with a mixing time of 50 ms to identify geminal LV methyl groups, and one with a mixing time of 300 ms to allow spin diffusion and buildup of a greater radius of NOEs. The pulse sequence for the ^1^H-^13^C HMQC was written in house with a t_1_ ^1^H refocusing composite pulse (63).

## Data, Materials, and Software Availability

Chemical shift assignments for all CM states described herein will be deposited in the BMRB prior to publication.

## Acknowledgments

This work was supported by NIGMS awards to Andrew L. Lee (GM127698 and GM144348). We would like to acknowledge Dr. Stuart Parnham at the UNC-Chapel Hill Biomolecular NMR Laboratory, which receives funding from the National Cancer Institute of the National Institutes of Health (P30CA016086). This study made use of NMRbox: National Center for Biomolecular NMR Data Processing and Analysis, a Biomedical Technology Research Resource (BTRR), which is supported by NIH grant P41GM111135 (NIGMS). We thank Dr. Kenneth Pearce of the Center for Integrative Chemical Biology and Drug Discovery for access to his PEAQ ITC instrument.

## Author Contributions

PJS and ALL designed research, PJS, TRM, and MSK performed research, PJS, TRM, and MSK analyzed data, KL and JA contributed reagents, PJS and ALL wrote the paper.

