## Supporting Information: Methods, Figures, Table for "Dynamic view of an allosteric intermediate in a positively cooperative dimer"

### Supporting Methods

#### *Synthesis of Transition State Inhibitor.*

All reactions were carried out in an oven-dried flasks under an inert argon atmosphere. Solvents and reagents were used as received unless otherwise noted. Diazomethylene)dibenzene was prepared according to the reported procedure.[1] Spectra for $^{1}H$ and $^{13}C$ NMR were recorded at room temperature with a Bruker (500 MHz and 126 MHz) with a dual carbon/proton cryoprobe. Chemical shifts are reported in δ (ppm) relative units to residual solvent peaks ($CDCl_3$ 7.26 ppm for $^{1}H$ and 77.16 ppm for $^{13}C$; $D_2O$ δ 4.79 ppm for $^{1}H$ NMR). Splitting patterns are assigned as s (singlet), d (doublet), t (triplet), q (quartet), quint (quintet), multiplet (m), dd (doublet of doublets), and td (triplet of doublets). Samples were analyzed on Waters Acquity UPLC with a PDA detector using C18 column, BEH 2.1 mm × 50 mm, 1.7 µm, at 30 °C with mobile phases A ($H_2O$ + 0.1% ammonium hydroxide) and B (MeCN). Mass spectrometry measurements were recorded using a Waters 3100 Mass Detector. Purification was carried out by an automated flash chromatography/medium-pressure liquid chromatography (MPLC). The diastereomers and enantiomers were analyzed on Agilent 1260 Infinity II using CHIRAPAK® IA or IC analytic column, 4.6 mm × 250 mm, 5 µm at 30 °C with mobile phases hexanes and isopropanol. The purification was carried out on TELEDYNE ISCO combiflash EZ PREP using CHIRAPAK® IA or IC semi-preparative column, 21 mm × 250 mm, 5 µm at rt with mobile phases hexanes and isopropanol.

**Synthesis of dibenzhydryl ester to resolve the endo enantiomer.**

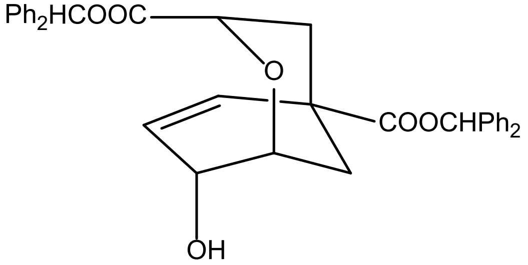


**(1*R*,3*S*,5*S*,8*R*)-1**

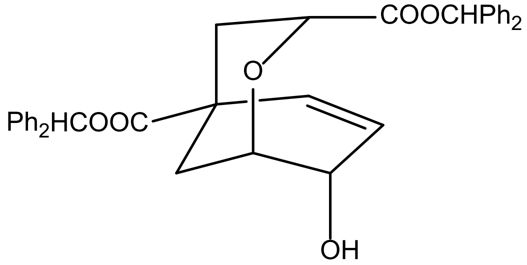


**(1*S*,3*R*,5*R*,8*S*)-1**

To a solution of 8-hydroxy-2-oxabicyclo[3.3.1]non-6-ene-3,5-dicarboxylic acid (3-endo-8-exo and 3-exo-8-exo mixture,104 mg, 458 µmol)[2-5] in MeOH (0.31 mL) and diethyl ether (2.8 mL) at 0 °C, was added (diazomethylene)dibenzene (622 mg, 3.2 mmol) until a yellow color persisted. The reaction was warmed to rt and quenched with AcOH until the color disappeared. Solvents were removed and the residue was purified via NP MPLC to give dibenzhydryl 8-hydroxy-2-oxabicyclo[3.3.1]non-6-ene-3,5-dicarboxylate (3-endo-8-exo and 3-exo-8-exo mixture, 29.2 mg, 11%) as a white solid.

The 3-endo-8-exo and 3-exo-8-exo mixture was separated using a semi-preparative CHIRALPAK® IA column to afford 22 mg of dibenzhydryl (3-endo-8-exo)-8-hydroxy-2-oxabicyclo[3.3.1]non-6-ene-3,5-dicarboxylate. The separation of enantiomers was performed using a semi-preparative CHIRALPAK® IC column eluted with 30% IPA/hexanes.

**(+)-Dibenzhydryl (1*R*,3*S*,5*S*,8*R*)-8-hydroxy-2-oxabicyclo[3.3.1]non-6-ene-3,5-dicarboxylate ((1*R*,3*S*,5*S*,8*R*)-1).** RT = 10.03 min, 10.5 mg, $\alpha_D^{20}$ = +24.5 (*c* = 0.42, DCM). $^{1}H$ NMR (600 MHz, $CDCl_3$) δ 7.37 – 7.28 (m, 20H), 6.89 (s, 1H), 6.88 (s, 1H), 5.75 – 5.70 (m, 1H), 5.26 (ddd, *J* = 10.0, 4.4, 1.4 Hz, 1H), 4.49 (dd, *J* = 7.7, 1.5 Hz, 1H), 4.30 (dd, *J* = 4.4, 2.2 Hz, 1H), 4.20 (dq, *J* = 4.0, 1.9 Hz, 1H), 2.63 (dt, *J* = 13.7, 2.1 Hz, 1H), 2.29 (dd, *J* = 13.6, 7.6 Hz, 1H), 2.08 (dt, *J* =

13.1, 2.2 Hz, 1H), 2.02 (ddt, $J$ = 13.0, 4.5, 1.2 Hz, 1H). $^{13}C$ NMR (151 MHz, $CDCl_3$) δ 173.15, 171.65, 139.96, 139.92, 139.60, 139.57, 131.39, 130.28, 128.78, 128.76, 128.68, 128.58, 128.33, 128.25, 128.22, 127.56, 127.51, 127.08, 127.04, 126.98, 77.76, 77.66, 72.81, 69.68, 65.19, 40.52, 31.87, 28.18. **(+)-1** obtained in this was assigned the structure shown from the X-ray structure.

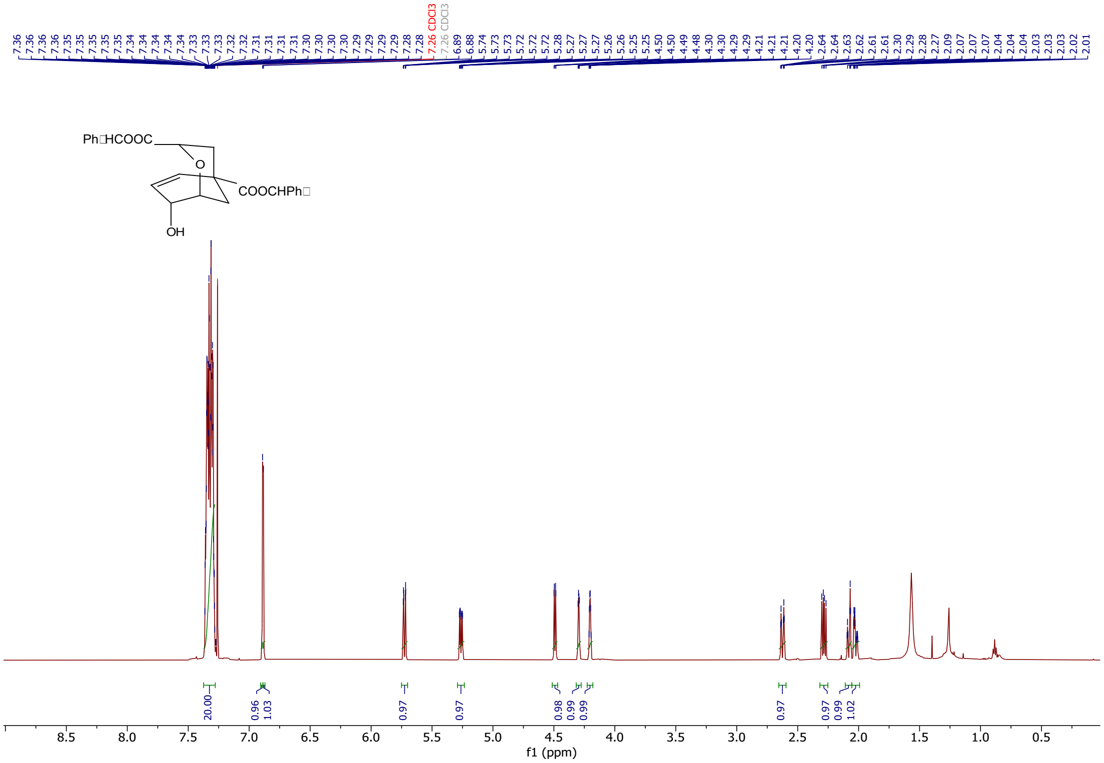


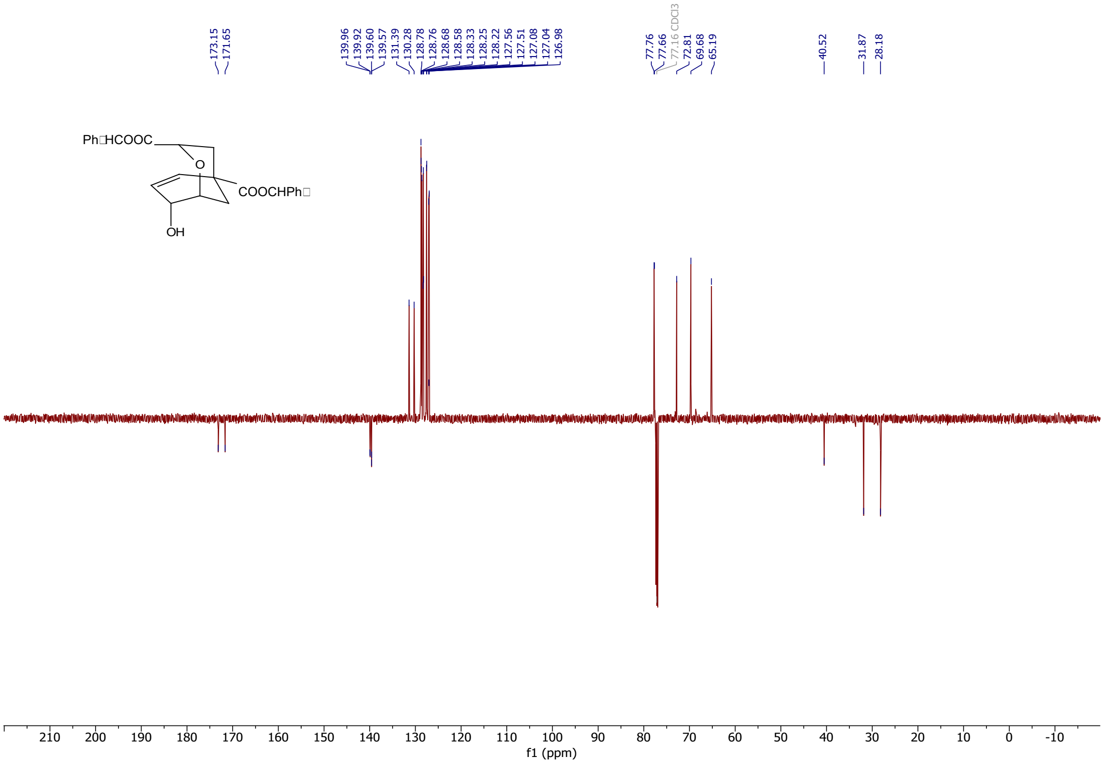

**(-)-Dibenzhydryl (1*S*,3*R*,5*R*,8*S*)-8-hydroxy-2-oxabicyclo[3.3.1]non-6-ene-3,5-dicarboxylate ((1*S*,3*R*,5*R*,8*S*)-1).** RT = 25.22 min, 10.4 mg. $\alpha_D^{20}$ = –27.4 (c = 0.42, DCM). $^1$H and $^{13}$C NMR spectra match those obtained for **(+)1**.

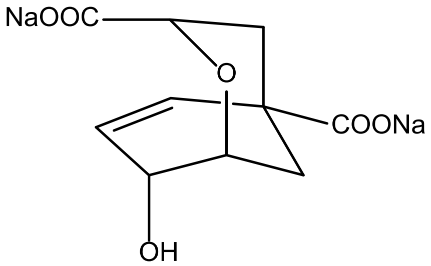

**(1*R*,3*S*,5*S*,8*R*)-2**

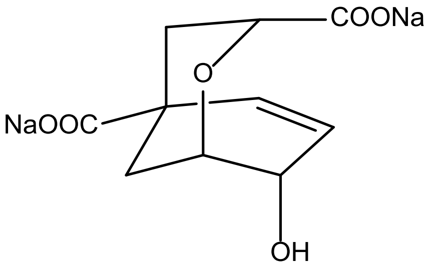

**(1*S*,3*R*,5*R*,8*S*)-2**

**Sodium (1*R*,3*S*,5*S*,8*R*)-8-hydroxy-2-oxabicyclo[3.3.1]non-6-ene-3,5-dicarboxylate ((1*R*,3*S*,5*S*,8*R*)-2).** To a solution of **(1*R*,3*S*,5*S*,8*R*)-1** (8.0 mg, 14 µmol) in MeOH (0.36 mL) and water (0.36 mL), was added NaOH (86 µL, 1.0 M, 86 µmol). The mixture was stirred at rt for 3 h. Solvents were removed and the residue was dissolved in water. The insoluble material was filtered and the filtrate was neutralized with 1N HCl. Water was removed; the residue was extracted with anhydrous MeOH. MeOH was removed and the residue was suspended in water, NaOH (29 µL, 1.0 M, 29 µmol) was added. Water was removed to afford the product, 4.9 mg, as a white solid and used without purification. $^1$H NMR (400 MHz, $D_2O$) δ 6.10 (d, *J* = 9.8 Hz, 1H), 5.76 (ddd, *J* = 10.0, 4.4, 1.5 Hz, 1H), 4.36–4.30 (m, 1H), 4.22 (dd, *J* = 7.2, 4.1 Hz, 1H), 4.21–4.10 (m, 1H), 2.23 (ddd, *J* = 13.6, 4.1, 1.8 Hz, 1H), 2.14 (dd, *J* = 13.6, 7.1 Hz, 1H), 2.03–1.89 (m, 2H).

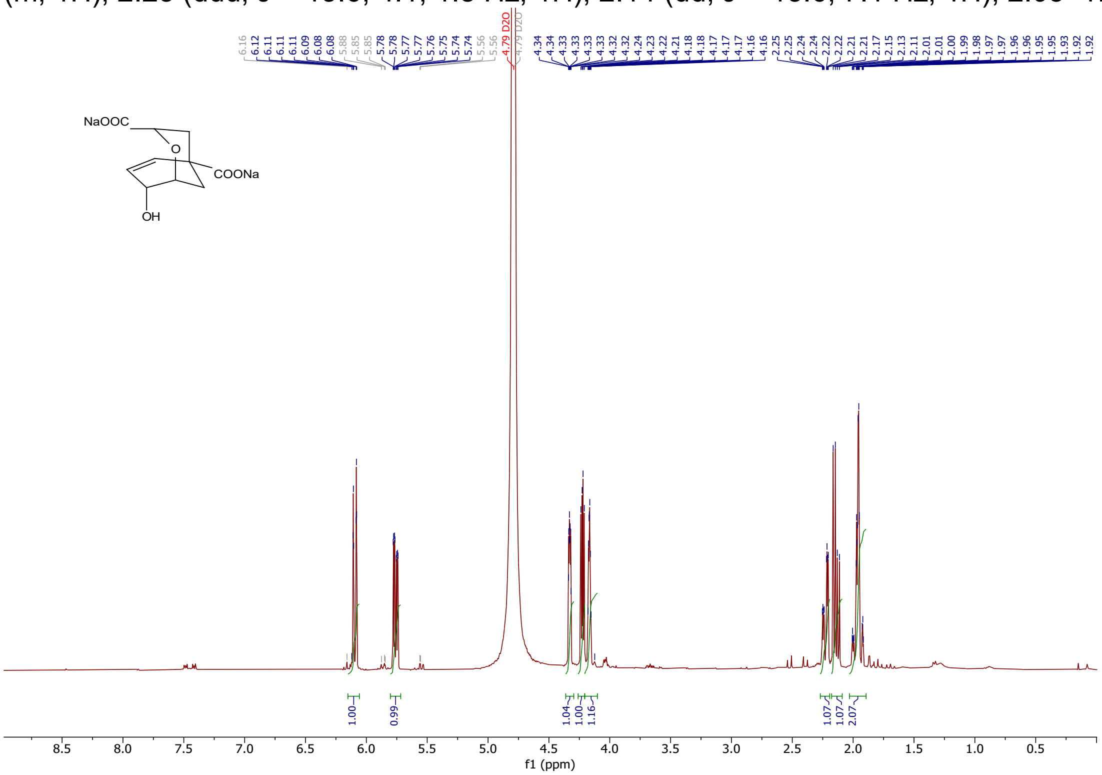

**Sodium (1*S*,3*R*,5*R*,8*S*)-8-hydroxy-2-oxabicyclo[3.3.1]non-6-ene-3,5-dicarboxylate ((1*S*,3*R*,5*R*,8*S*)-2).** To a solution of **(1*S*,3*R*,5*R*,8*S*)-1** (8.0 mg, 14 µmol) in MeOH (0.36 mL) and water (0.36 mL), was added NaOH (86 µL, 1.0 M, 86 µmol). The mixture was kept at rt for 3 h. Solvents were removed and the residue was dissolved in water. The insoluble material was filtered and the filtrate was neutralized with 1N HCl. Water was removed; the residue was extracted with anhydrous MeOH. MeOH was removed and the residue was suspended in water, NaOH (29 µL, 1.0 M, 29 µmol) was added. Water was removed to afford the product, 6.1 mg, as a white solid and used without purification. $^{1}H$ NMR spectra match that obtained for **(1*R*,3*S*,5*S*,8*R*)-2**.

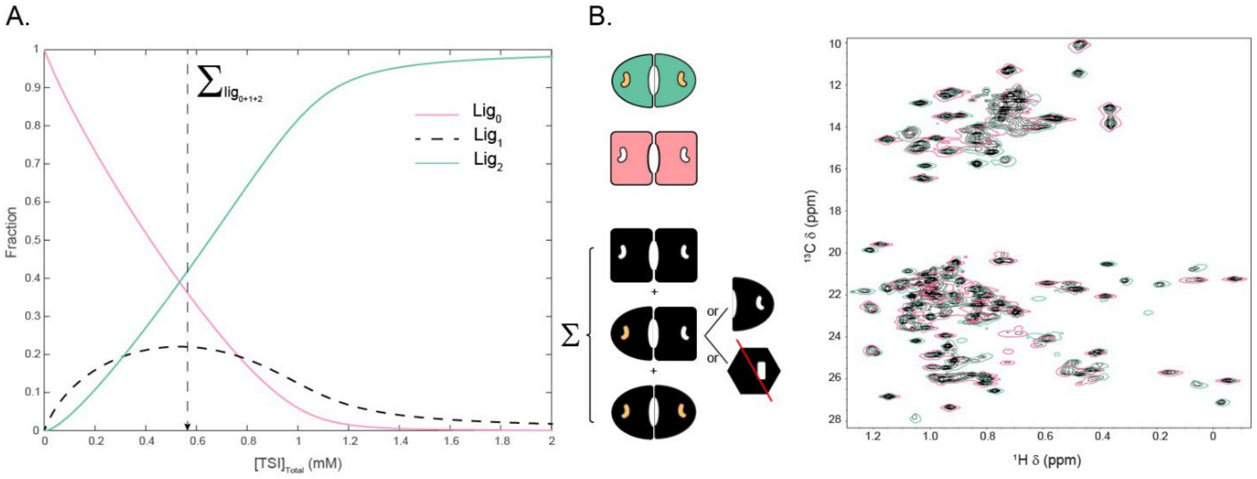


***Figure S1.*** (A) Fraction species for TSI binding to CM based on a two-site binding polynomial parameterized by the ITC data in Figure 1. The population of $lig_1$ reaches a maximum of only 20% in this case due to positive cooperativity. (B) ILV HMQC at peak [$lig_1$] conditions—vertical dotted line in panel A— is shown in black and compared with the symmetrical $lig_0$ (T-state structure) and $lig_2$ (superR structure) endpoint spectra in single contours. Note that black $lig_1$ resonances only show up under $lig_0$ and $lig_2$ peaks indicating subunits of the $lig_1$ state must resemble the T or R conformations or be broadened to the point of invisibility at a population of 20%.

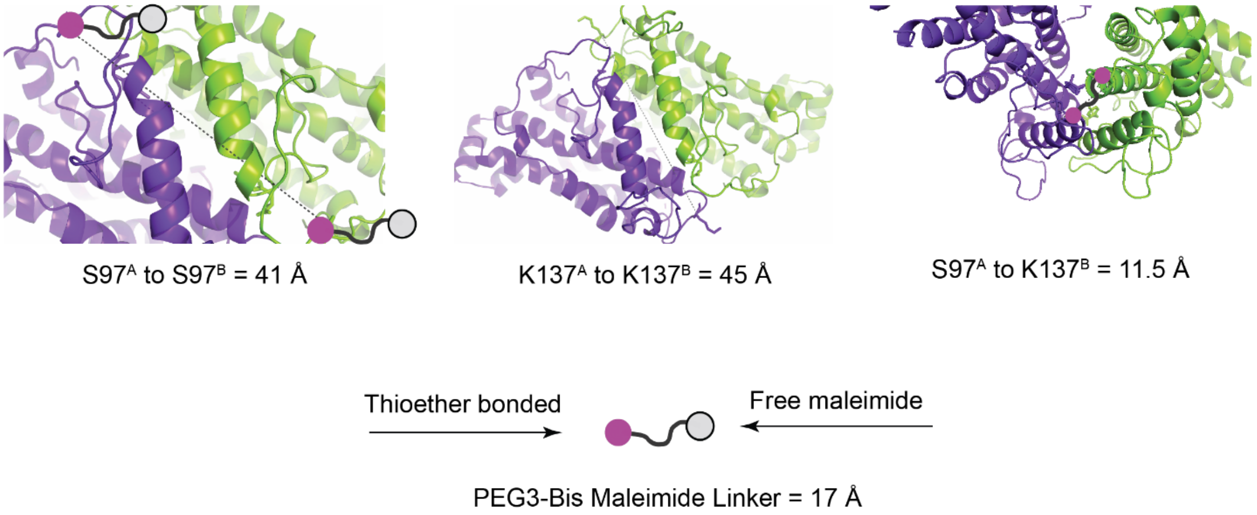


***Figure S2.*** Geometry of residues chosen for linkage amino enables stepwise tagging then linkage, and hence a near quantitative yield of desired MLD. Tagging the first dimer alone (C97 for example) reacts with all exposed cysteines attaching a *distinct* Bis Mal linker to both C97 residues in the dimer; no linkage via the second maleimide is possible due to the distance of the two cysteines in the dimer relative to the length of the linker. However, after removal of excess linker from the tagging reaction and upon adding the second dimer, the second cysteine is brought into reactive proximity of the exposed, unreacted second maleimide head.

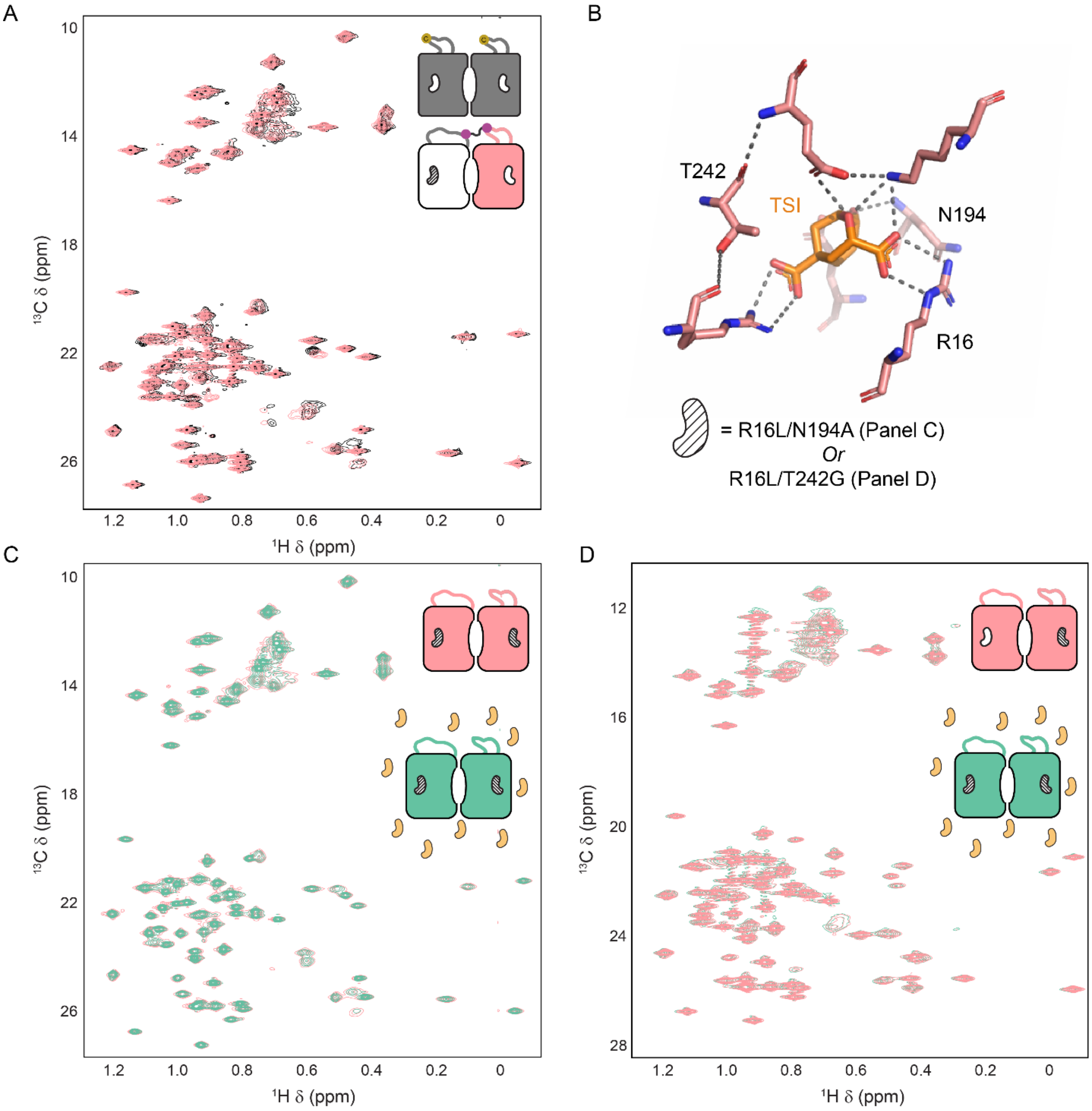


***Figure S3.*** (A) Linkage does not affect the structure which is consistent with retention of critical activity and regulatory features shown in Figure 2B of the main text. (B) CM active site residues that coordinate the TSI are shown in sticks. Two pairs of mutants were made to abolish substrate binding: R16L/N194A and R16L/T242G. Neither has residual catalytic activity (data not shown), and neither binds a trace of TSI under the conditions used to generate $lig_1$ dimers (C and D). In spectra of both double mutants, resonances of samples with and without TSI are perfectly overlapped. In addition, the spectra with TSI are not broadened at all, ruling out low affinity binding leading to bi-molecular $R_{ex}$ as the origen of line broadening in $lig_{1E}$.

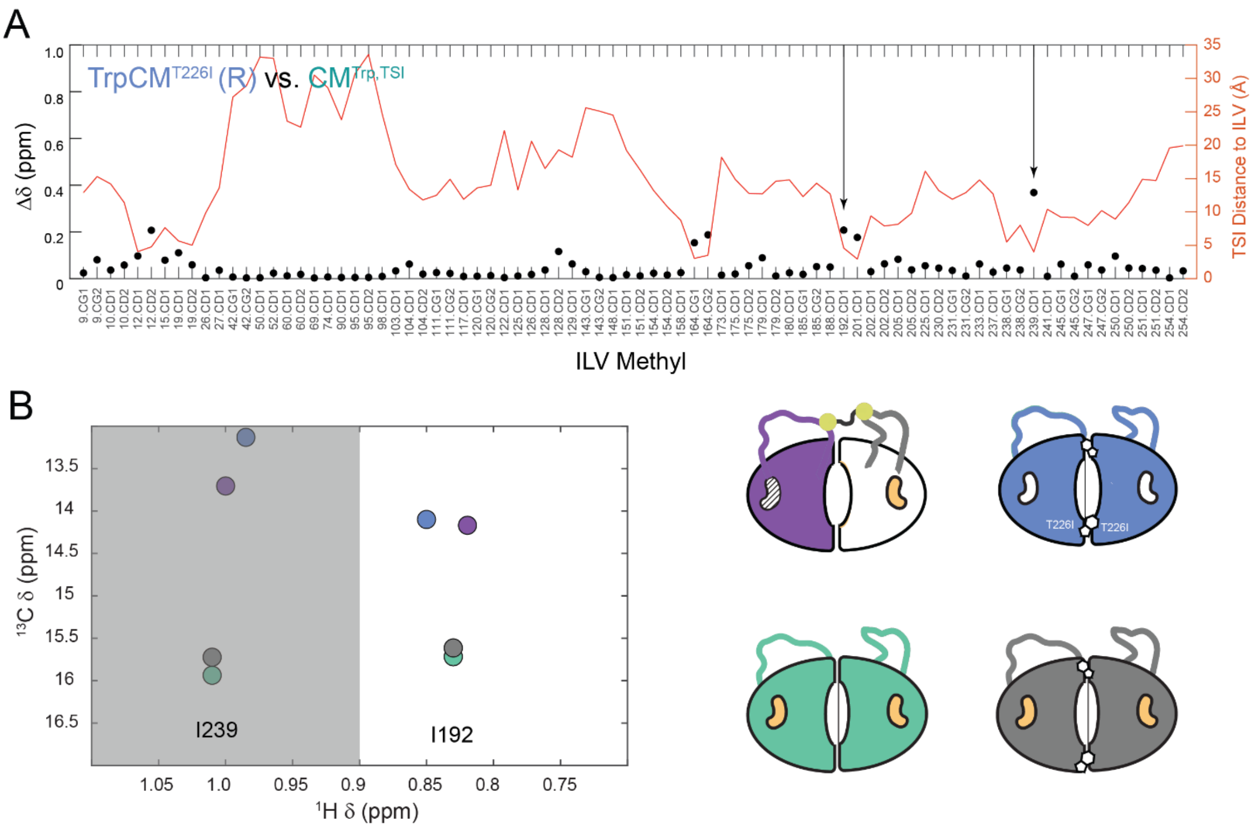


***Figure S4.*** Chemical shifts of diagnostic active sites residues argue against residual binding in $lig_1$ mutated active sites. Although we showed TSI does not bind the R16L/N194A or R16L/T242G mutant active sites in a homo dimer setting (Figure S3C&D above), we considered that positive cooperativity could render the second, mutant active site of a $lig_1$ MLD higher in affinity than what is measured in the control experiments (Figure S3C&D). To rule out any binding in the mutant active sites in $lig_1$, we identified a few resonances that are different in the R-free vs superR-active site bound forms (I239 and I192, arrows in panel A) and asked if the $lig_{1E}$ resonances clustered with the empty or bound control peaks (panel B). The purple $lig_{1E}$ resonances of the MLD clearly map to the active site empty R-state, effectively ruling out unwanted binding.

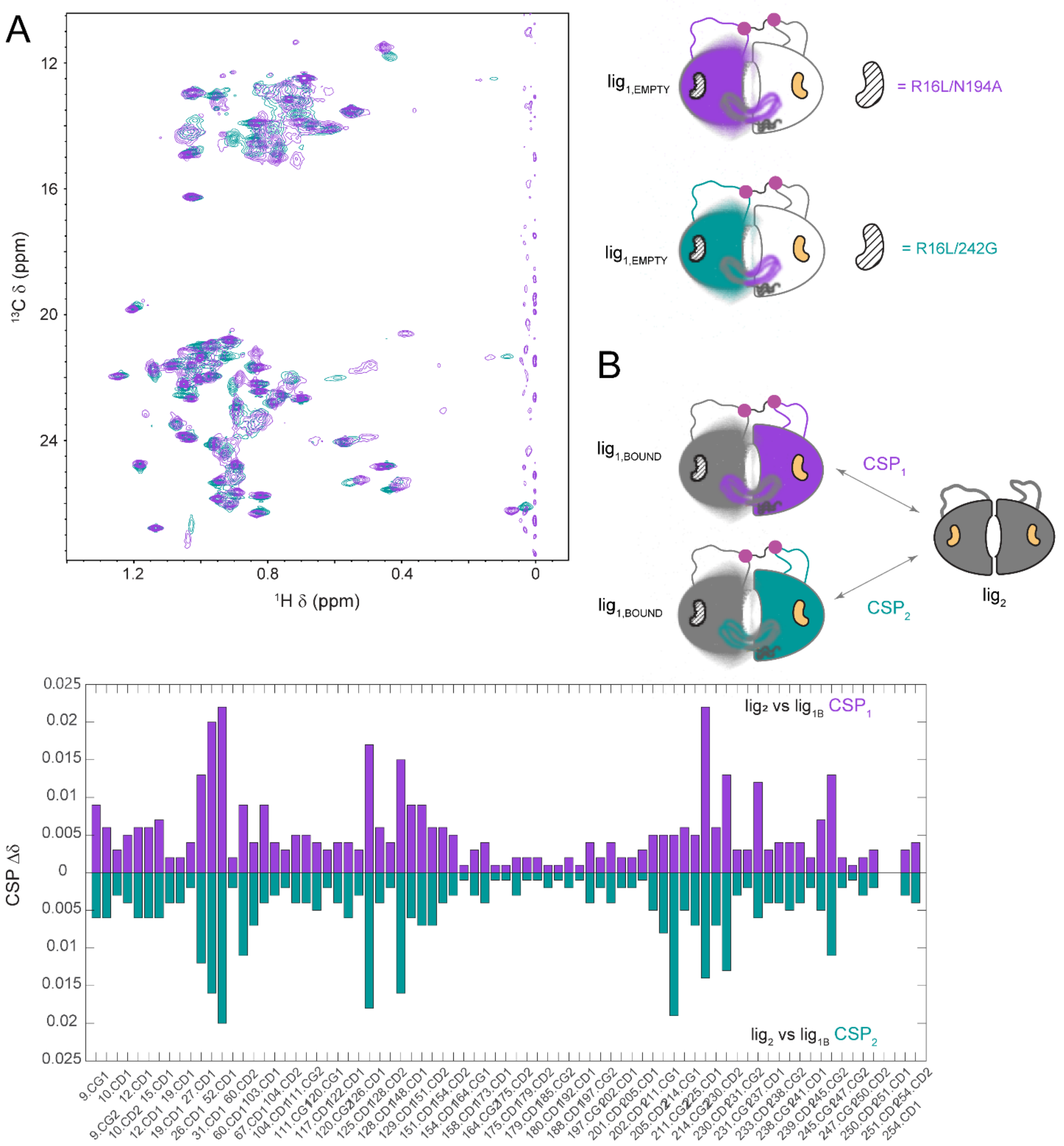


***Figure S5.*** Similar chemical shift and dynamic behavior in MLDs with two different active site mutant pairs. (A) Lig$_{1E}$ ILV HMQC spectra for the R16L/N194A (purple) and R16L/T242G (cyan) MLDs have excellent overlap indicating both are in the R-state (see in depth chemical shift comparison in Figure 3 of the main text). The broadening phenomenon is apparent in both active site pairs, reinforcing the conclusion that the empty subunit of lig$_{1E}$ is fuzzy. (B) The differences in lig$_{1B}$ vs lig$_2$ are subtle but mirrored between two independent active site mutation pairs, and therefore represent meaningful effects of second ligand binding to the pre-bound active site. These effects contribute to cooperativity ($\Delta G_1$ vs $\Delta G_2$).

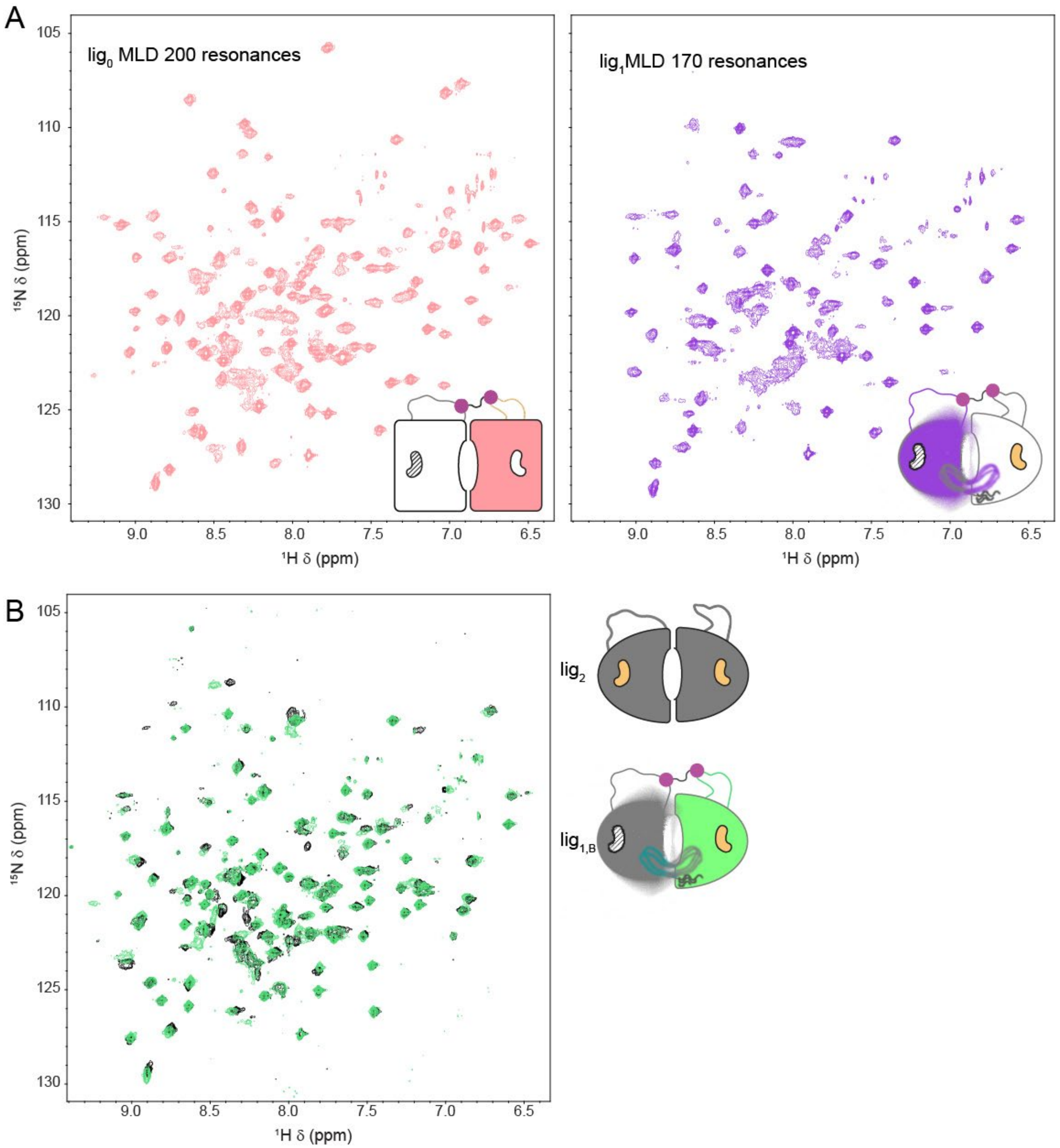


***Figure S6.*** Backbone perspective of the lig$_1$ complex reinforces conclusions based on side-chain ILV resonances. A) Comparison of the lig$_0$ and lig$_{1E}$ MLD showing dramatic peak shifting to an R-conformation and significant line broadening evidenced by a complete loss of 30 peaks. (B) Near perfect overlap of the lig$_{1B}$ (green) and lig$_2$ spectra indicating a switch to a more canonical super-R conformation in the local subunit, but with subtle differences that are described in detail in Figure 5 and the main text.

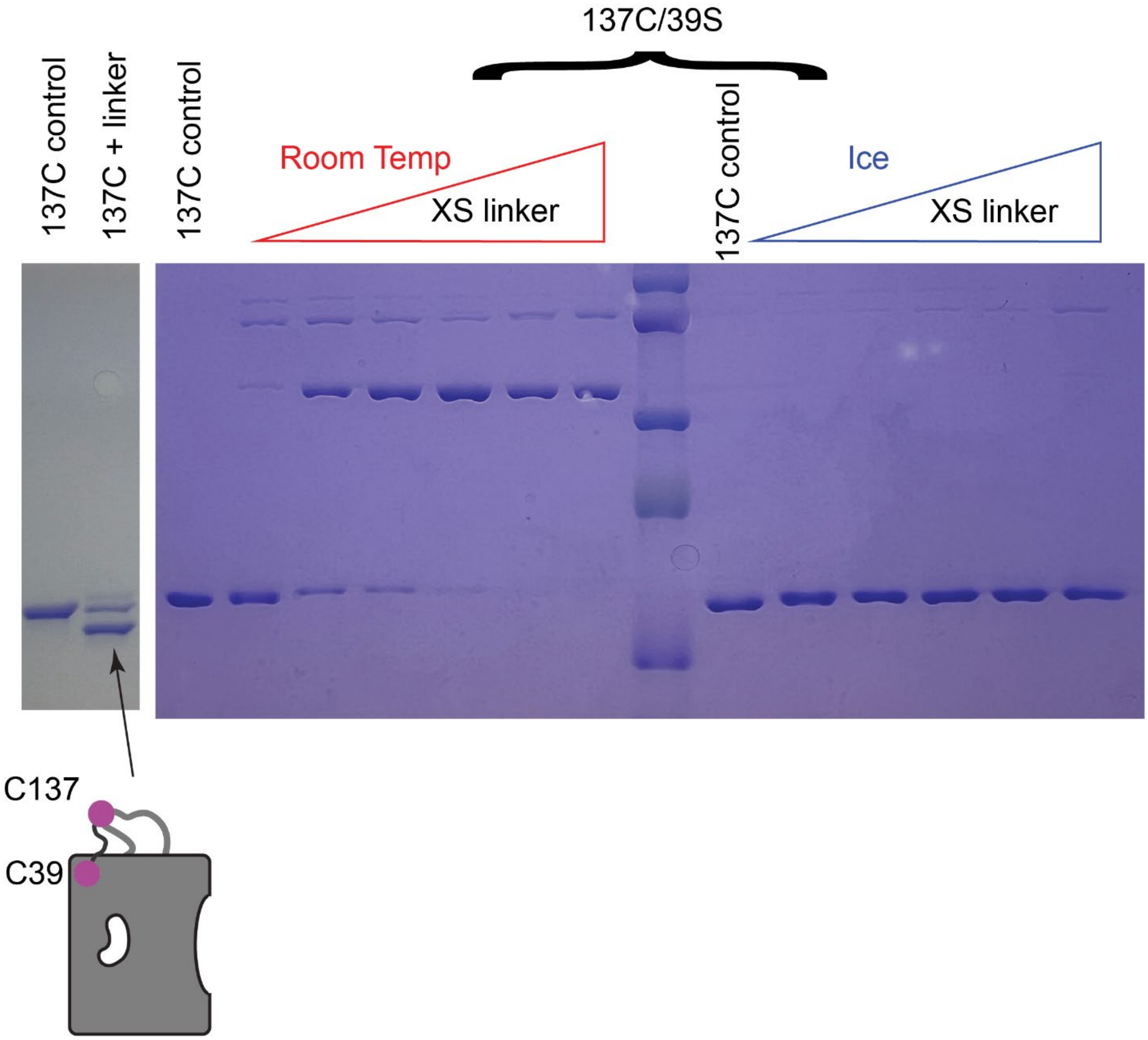


***Figure S7.*** Attempts to tag C137 with bis-mal-PEG3 in a wild-type background resulted in some intra-protomer linkage with a different exposed cysteine (Left). We deduced it was C39 based on proximity, and the intra protomer linkage was prevented in a C39S background (see minor band migration retardation in the 'ice' set of tagging reactions in the second gel. The second gel also shows that tagging the first dimer on ice was required to prevent trans dimer, C137 to C137 linkage, which should not be possible given the 137 to 137 distance across the dimer interface and length of the linker (see Figure S2 above).

**Table S1.** MLDs used in this study

| MLD | WT Active Site Subunit | Mutant Active Site Subunit | Linkage Pair |
|---|---|---|---|
| Lig1E-1 | C39S/K137C | *R16L/N194A/S97C | 97X137 |
| Lig1E-2 | C39S/K137C | *R16L/T242G/S97C | 97X137 |
| Lig1B-1 | *C39S/K137C | R16L/N194A/S97C | 97X137 |
| Lig1B-2 | *C39S/K137C | R16L/T242G/S97C | 97X137 |

* Isotopically labeled subunit
